# Tropomodulin and Leiomodin Differentially Nucleate Tropomyosin Assembly on Actin Filaments

**DOI:** 10.64898/2026.10.04.756620

**Authors:** Camille Bagès, Dina Al Abyad, Rebecca Pages, Shayna B. Brotzman, Thomas Dos Santos, Guillaume Romet-Lemonne, Roberto Dominguez, Antoine Jégou, Hugo Wioland

## Abstract

Tropomyosins play a central role in the regulation of the actin cytoskeleton by assembling along filaments into rope-like, head-to-tail polymers (“clusters”) along filaments that regulate the binding and activity of numerous actin-binding proteins. Importantly, in cells, tropomyosins do not decorate actin filaments uniformly; some filaments are devoid of tropomyosins, whereas others recruit specific isoforms. To explore the diversity of tropomyosin-isoform binding kinetics and identify factors that control their assembly, we designed tropomyosin constructs, bearing a natural N-terminal acetylation and targeted cysteine mutations for fluorescence labelling. We analysed the binding of different tropomyosin isoforms to individual actin filaments reconstituted inside a microfluidic chamber. We show that, despite significant differences in spontaneous nucleation and elongation rates, all tested tropomyosin isoforms can coassemble within the same clusters. In addition, we identified tropomodulin and leiomodin, two pointed-end-binding proteins, to efficiently nucleate the formation of tropomyosin clusters on actin filaments, for all tested tropomyosin isoforms. Notably, while tropomodulin nucleates tropomyosin clusters from the pointed end, leiomodin does so along the entire filament.

## Introduction

In mammalian cells, actin assembles into a wide array of filament networks, each characterized by a unique protein composition, architecture and dynamics, which ultimately determine the function of each network [1], [2]. With the exception of muscle sarcomeres, these filament networks are essentially composed of two actin isoforms: cytoplasmic β- and γ-actin. β- and γ-actin are nearly identical, differing by only four amino acid substitutions [3]. As a consequence, how actin filament and network diversity emerges has long been a central focus of cytoskeleton research [1], [2].

Tropomyosins (Tpm) are a key family of proteins that have been proposed to determine actin filament (F-actin) identity [1], [4]. In humans, Tpm are encoded by four genes resulting in dozens of isoforms through alternative splicing [5]. In cells, actin networks show large disparities with some being almost devoid of Tpm (e.g. lamellipodia, filopodia) while others concentrate most of the Tpm (typically contractile networks such as stress fibers) [6]. In addition, Tpm isoforms show preferences for specific actin networks, and they often do not colocalize [6]. Once bound to F-actin, Tpm regulate the binding and activity of other actin-binding proteins in an isoform-specific manner [4], [7], [8], [9], [10], [11]. Tpm isoforms thus help define the identity and function of the F-actin they decorate. Understanding how, when, and where Tpm decorate filaments is therefore essential to understanding actin network diversity.

At the molecular level, Tpm form parallel coiled-coil dimers that bind cooperatively head-to-tail along the long-pitch helix (strand) of the actin filament, forming polymer-like structures, here called clusters (Fig. 1A). Each Tpm dimer covers 6 or 7 actin subunits, depending on the Tpm isoform. However, single Tpm dimers have weak affinity for F-actin [12], and it is through the N- to C-terminal cooperative association of consecutive dimers that Tpm acquire measurable affinity for F-actin, leading to the formation of clusters on the filament. Once the step of nucleation (i.e. the assembly of a minimal stable Tpm cluster) is complete, Tpm clusters elongate and disassemble through the addition and dissociation of Tpm dimers at the two ends of the clusters (Fig. 1A) [7], [11], [12]. While the cooperative binding mechanism is qualitatively conserved among Tpm isoforms, affinity and cooperativity vary widely [10], [13]. Interestingly, some isoforms have been reported to coassemble within the same clusters, while others compete for binding [8]. Yet, fully characterising the assembly of different Tpm isoforms remains challenging, notably to purify functional constructs (Fig. 2) [13], [14].

**Figure 1:**
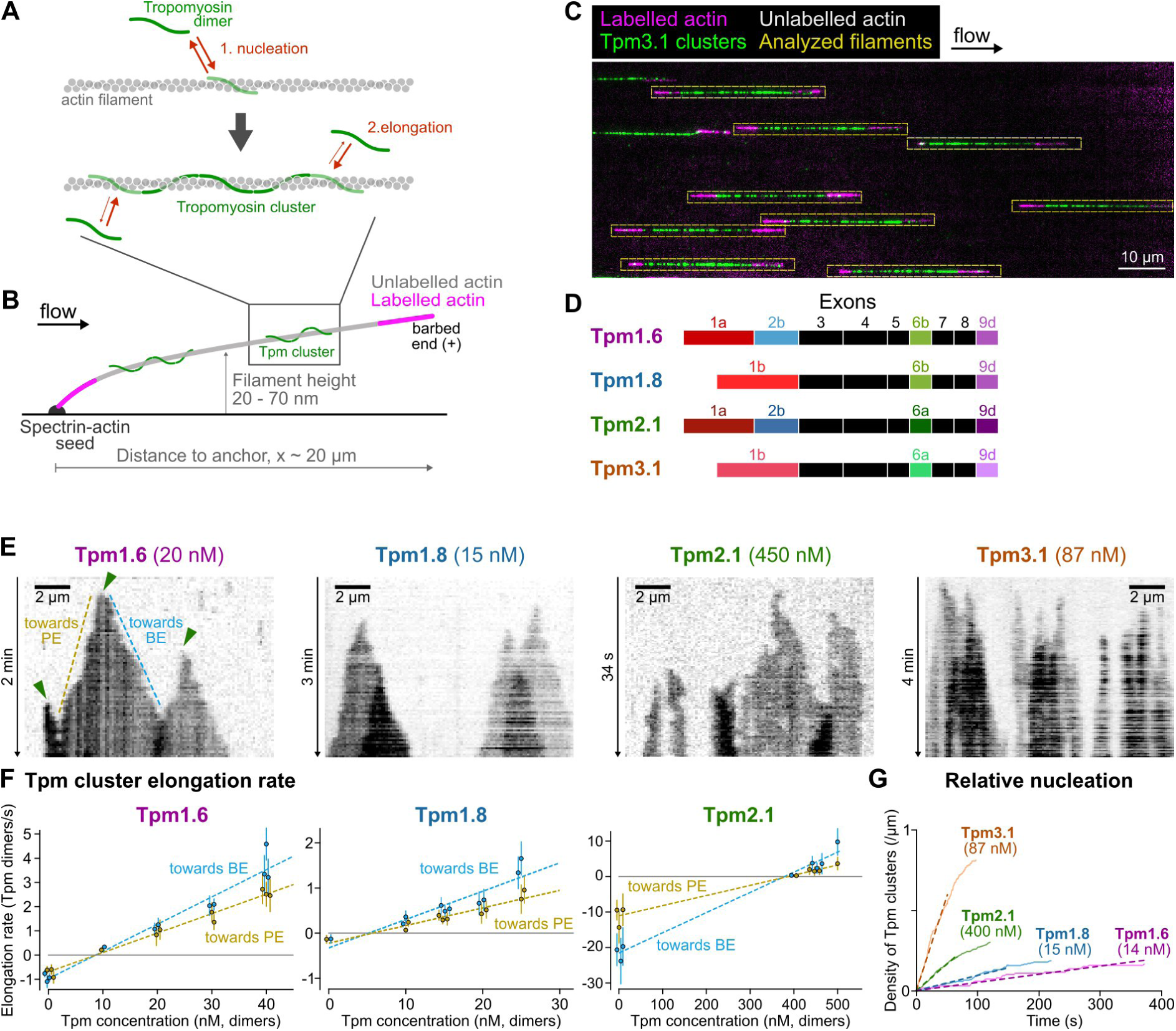
Differential binding dynamics of Tpm isoforms. **A.** Tpm clusters form in two steps: nucleation and elongation towards the actin filament ends. **B.** Experimental setup. Single actin filaments are polymerized from surface-anchored spectrin-actin seeds, with a long unlabelled segment flanked by two short fluorescent segments. Tpm clusters assembly is only quantified on the unlabelled segment. **C.** Partial view of a typical field of view. Tpm3.1 partially cover unlabelled segments of F-actin. **D.** Exon composition of the four tested Tpm isoforms. Coloured exons correspond to alternatively spliced ones. **E.** Typical kymographs of the binding of different Tpm isoforms. Note the different time scales and Tpm concentrations. Arrow heads indicate nucleation events and dashed lines highlight the elongation towards the filament barbed end (BE) and pointed end (PE). The BE is always on the right hand side of kymographs. **F.** Quantification of the elongation and disassembly rates of Tpm clusters towards the two filament ends. Individual points correspond to averages over a one experiment (N =7 to 34 filaments). Error bars correspond to standard deviations. Note that for readability, data points at the same concentration are displayed as swarmplots not to overlap. We were not able to measure the elongation rate for Tpm3.1 because of its very high nucleation rate. **G.** Relative nucleation to elongation rates of the four isoforms. The Tpm concentrations were adjusted such that clusters elongate at about the same rate (Fig. S1D). N = 12 to 17 filaments per condition. Dashed lines: linear fits. Tpm3.1 nucleates 3.5x faster than Tpm2.1; Tpm2.1 3.4x faster than Tpm1.8; and Tpm1.8 2x faster than Tpm1.6.

**Figure 2:**
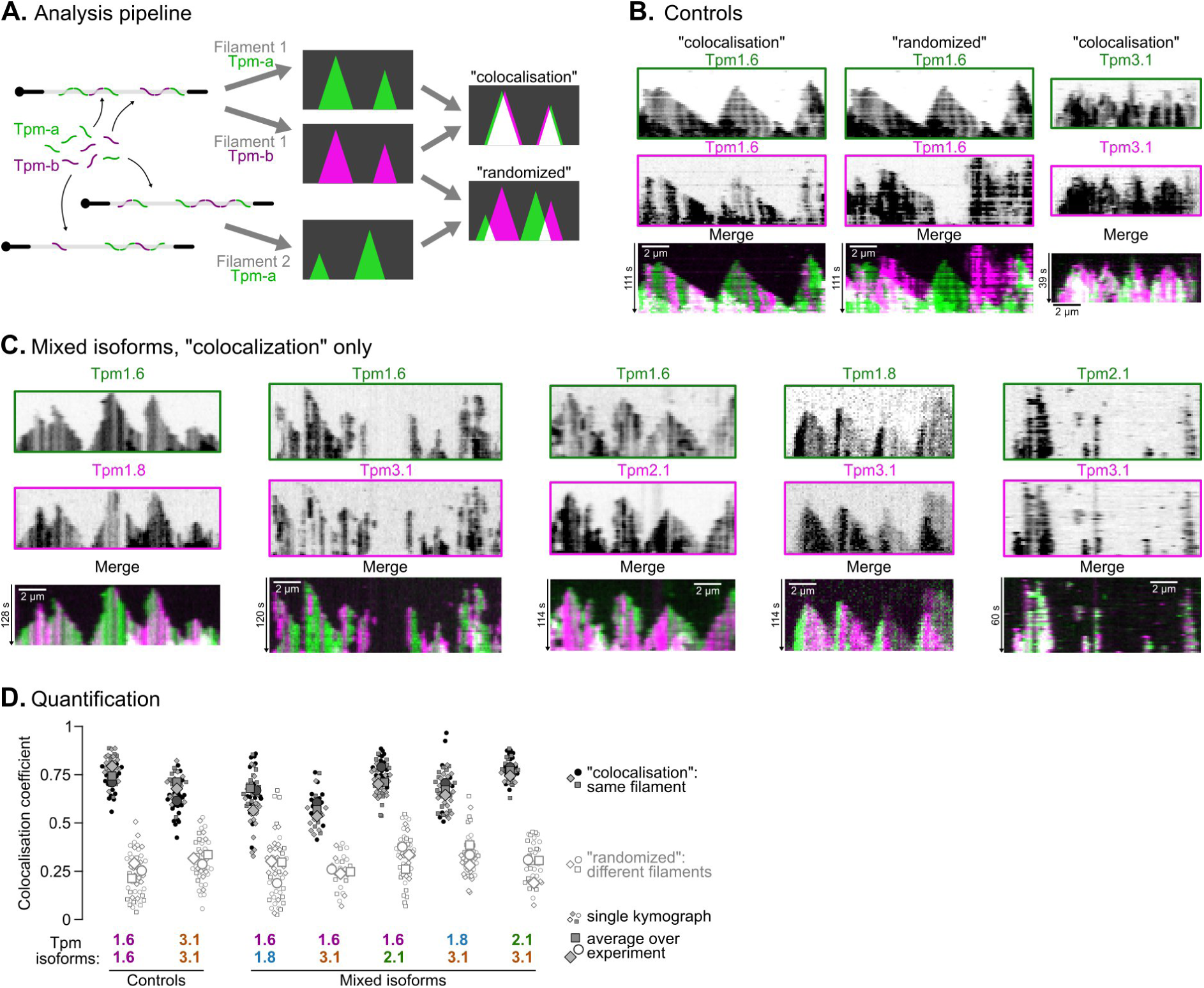
Coassembly of Tpm isoforms. **A.** Analysis pipeline for microfluidics Tpm isoform colocalization assays. Single actin filaments are exposed to a mix of Tpm labelled with different fluorophores. Kymographs are constructed either by taking the two channels from the same filament, to measure the actual colocalisation coefficient, or from two different filaments as a negative control. **B-C.** Typical coassembly kymographs obtained for either the same (B) or different Tpm isoforms (C). All kymographs start from t=0 when filaments are exposed to Tpm. Same scale bar for all kymographs. **D.** Colocalisation coefficient obtained for all combinations of Tpm isoforms.

The binding of Tpm to F-actin has been proposed to be regulated by actin filament nucleators and elongators [1], [15]. Experiments in the yeast *S. pombe* showed that the formins For3 and Cdc12 determine the localisation of specific Tpm variants (acetylated and unacetylated Cdc8) [16]. However, in vitro experiments using purified yeast and mammalian proteins have so far failed to demonstrate direct or specific recruitment of Tpm by formins [15], [17]. More generally, no actin-binding protein has been shown so far to directly promote Tpm assembly in vitro, to our knowledge. Here, we reveal that the protein family Tropomodulin/Leiomodin (Tmod/Lmod) could be one of those players.

There are four Tmod and three Lmod human isoforms, showing both similarities and differences at the sequence level, which correlate with their different biochemical and cellular activities [18], [19]. Tmod features two Tpm- and two actin-binding domains (TpmBS1–ABS1–TpmBS2–ABS2 (LRR), Fig. 3A), which together mediate tight pointed end (PE) binding, through interactions with the first three actin subunits as well as with the two Tpm dimers on each filament strand (Fig. S3A) [20], [21], [22]. In this way, Tmod acts as a capping protein, inhibiting subunit exchange at the PE, and this activity is greatly enhanced through interaction with Tpm. In cells, they are responsible for the stability of a number of actin networks, such as sarcomeres, stress fibers, and spectrin-actin networks [19], [23]. The depletion of Tmod in cells induces the disassembly of Tpm-decorated F-actin [18], [23]. Because the affinity of Tmod for the PE is largely enhanced by Tpm, it is thought that Tpm first decorate actin filaments and then help recruit Tmod to the PE [19]. Here, we propose an alternative hypothesis: Tmod initiates the assembly of Tpm clusters at filament PE.

**Figure 3:**
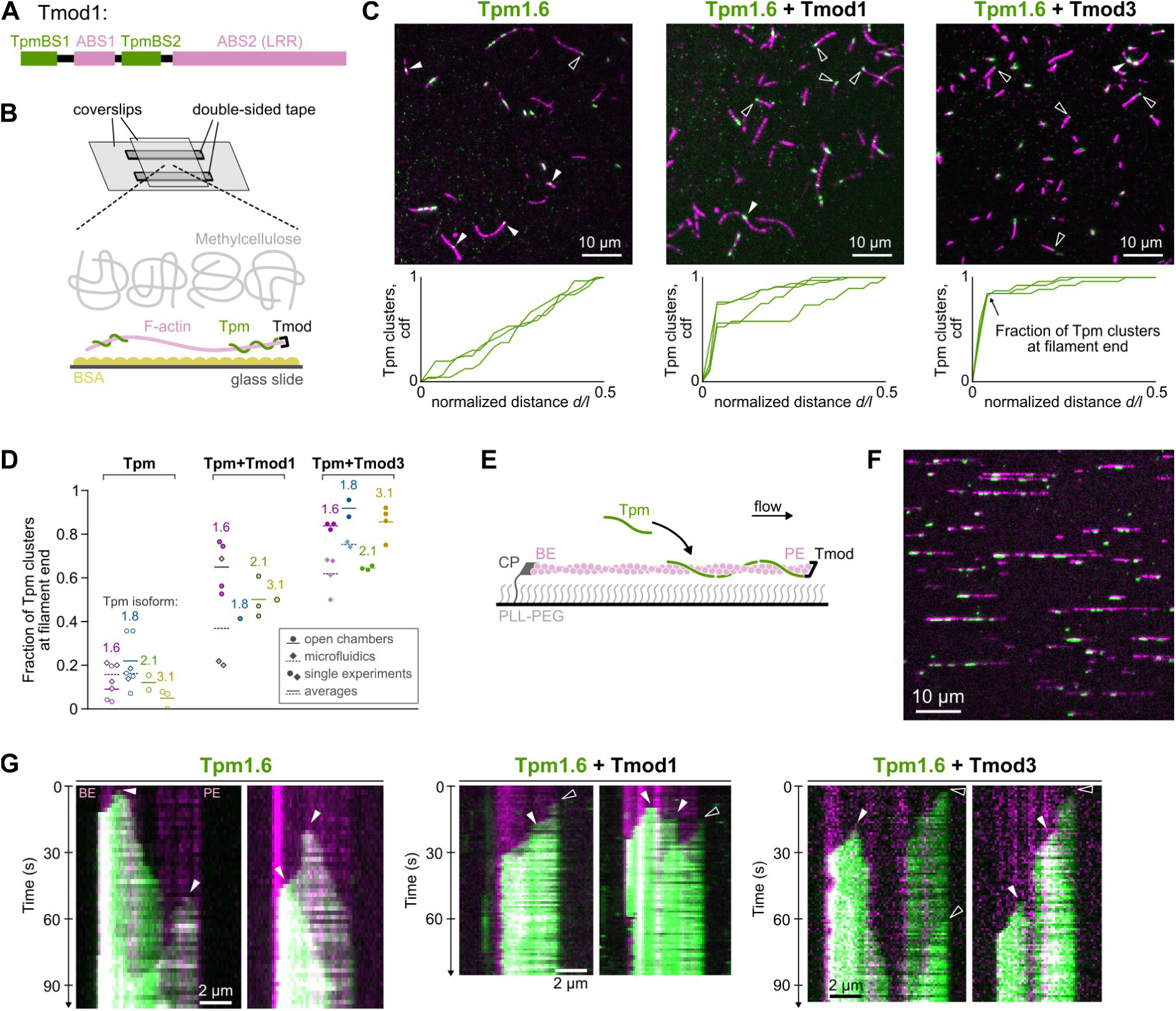
Nucleation of Tpm clusters by Tmod in vitro. **A.** Domain structure of tropomodulins. TpmBS: tropomyosin binding site, ABS: actin bindinding site, LRR: leucine rich region. **B.** Open chamber setup. Single actin filaments are confined near the bottom coverslip by methylcellulose. **C.** Biased localisation of Tpm1.6 by Tmod in open chambers. Top: Typical snapshots of the experiment, taken 1 to 2 min after mixing single actin filaments (magenta), 20 nM Tpm1.6 (green), ± 50 nM Tmod. The delay after mixing was chosen such that there was one Tpm cluster per actin filament, on average. Black and white arrowheads highlight Tpm clusters close to or away from the filament end, respectively. Bottom: cumulative distribution functions (cdf) of the position of Tpm clusters to the filament closest end, *d/l*. Each line corresponds to a single experiment. N = 16-61 filaments per experiment (mean: 32). **D.** Tmod increases the fraction of Tpm clusters first nucleated at the filament pointed end. Individual symbols: value of the cdf at *d/l = 0.04*. Round and square symbols: experiments performed in open and microfluidic chambers, respectively. Colours: Tpm isoform. Averages are calculated for open and microfluidic chambers separately. N=14 to 78 filaments per experiment (average: 33). Filament length *l* in open chambers: 3.5-9.5 µm (mean: 6.9 µm). In microfluidics, 5.9 µm segments starting from the free pointed end where analysed. See supplementary data for full statistics. **E.** Microfluidic setup with BE anchoring by CP, exposing the PE to Tmod. **F.** Part of a typical field of view with Tpm1.6 (green) bound to single actin filaments (magenta). **G.** Typical kymograph showing the nucleation (arrowheads) and elongation of Tpm1.6 clusters. The elongation shows irregularities and interruptions which we attribute to the labelling of actin filaments (5% Alexa-568, magenta).

Lmod share with Tmod TpmBS1 and the LRR domain (ABS2), but lack the second Tpm-binding site and most of ABS1, and contain a C-terminal extension featuring a Proline-rich domain (PRD) and an actin-binding WASP-Homology 2 (WH2) domain (Fig. 4A) [18], [19]. These differences in domain architecture correlate with differences in activity between the two subfamilies, with Lmod displaying both nucleation [19] and PE elongation [24], [25] activities but not capping activity. Moreover, contrary to Tmod that shows sharp localization to two thin bands on each side from the M-line in muscle sarcomeres, Lmod shows more diffuse localisation, being enriched at PEs but also distributed along the length of thin filaments. This localisation could in principle allow Lmod to help localise Tpm to filament sides.

**Figure 4:**
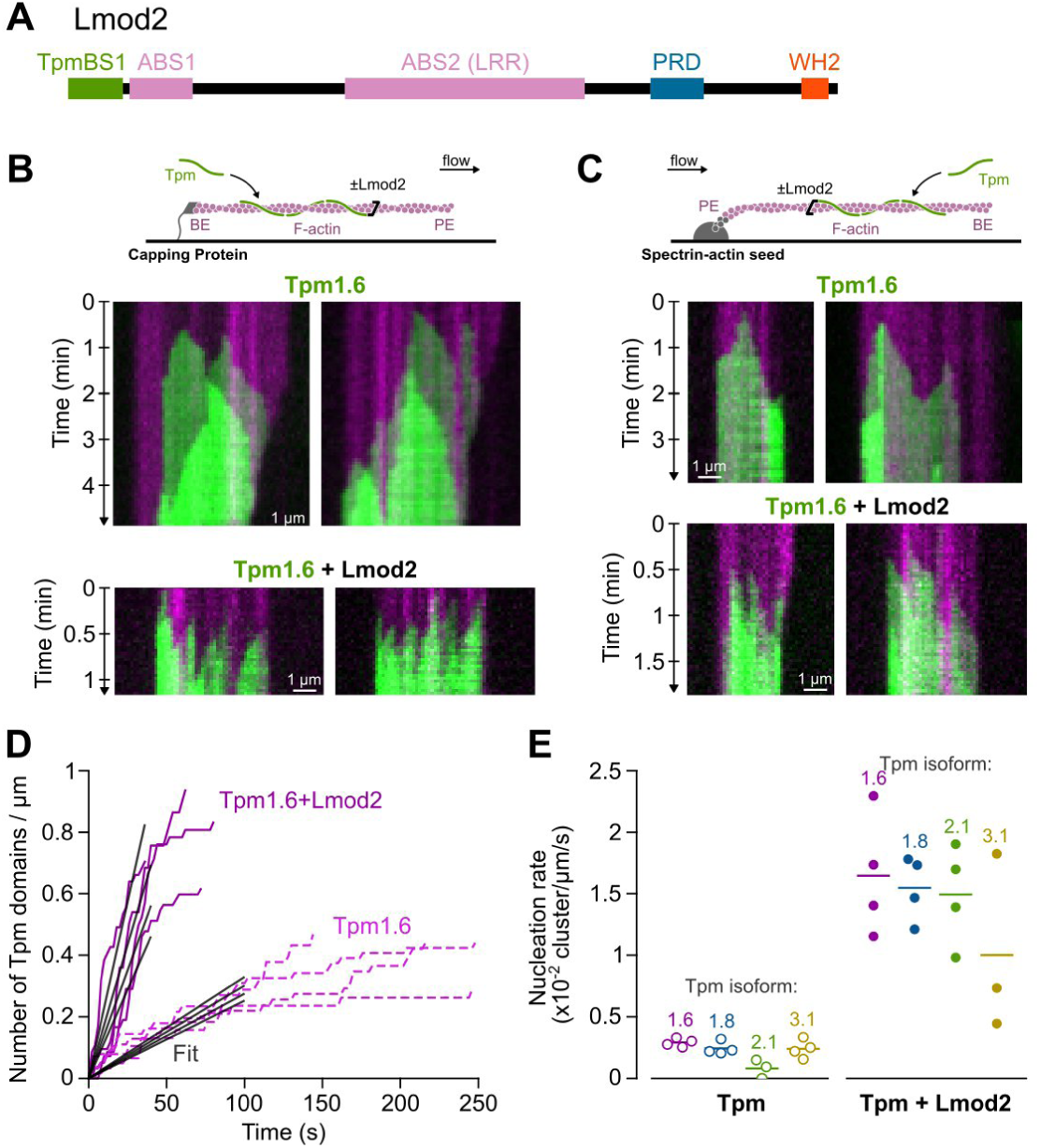
Nucleation of Tpm clusters by Lmod2 in vitro. **A.** Domain structure of Lmod2. TpmBS: tropomyosin binding site, ABS: actin bindinding site, LRR: leucine rich region, PRD: proline-rich domain, WH2: WASP-Homology 2 domain. **B-C.** Tpm1.6 cluster nucleation and assembly over Capping Protein (A) and spectrin-actin seed (B) anchored actin filaments. Filaments were exposed from time *t=0* to 10 nM Tpm1.6 ± 50 nM Lmod2. **C.** Quantification of the density of Tpm1.6 clusters over time. Each line corresponds to a single experiment. N = 5-8 filaments, 11-36 clusters. Linear fits (grey lines) were used to calculate the nucleation rate. **D.** Lmod2 increases the nucleation rate for all tested Tpm isoforms. Dots: single experiments. Lines: averages. N = 4-10 filaments, 11-56 clusters per experiment.

Here, we address the binding of Tpm isoforms to actin filaments and its regulation by Tmod/Lmod-family proteins. We quantify the dynamics of Tpm recruitment to single actin filaments, reconstituted inside a microfluidic chamber. We compare the assembly kinetics of four different human Tpm isoforms, using constructs featuring natural N-terminal acetylation and fluorescence labelling at cysteine residues introduced by mutagenesis (Fig. 1). We show that all tested Tpm isoforms are capable of coassembling (Fig. 2). We further reveal that both Tmod and Lmod are potent nucleators of Tpm cluster assembly, with Tmod nucleating Tpm clusters at the filament PE while Lmod nucleates Tpm along the filament length (Fig. 3-4).

## Results

### 1. Tpm isoform exhibit quantitatively distinct binding kynetics

While human cells can express dozens of Tpm isoforms through alternative splicing, this study focuses on four cytoplasmic isoforms - Tpm1.6, Tpm1.8, Tpm2.1 and Tpm3.1 - which represent three different genes, long and short isoforms [6] (Fig. 1D). These isoforms have been largely studied both in vitro and in cellulo. In cells, Tpm isoforms are often associated with fiber-like networks, such as stress fibers and transverse arcs. As a consequence, Tpm1.6, Tpm2.1 and Tpm3.1 are all necessary for proper stress fiber formation by promoting myosin 2 activity and protecting filaments from cofilin-mediated disassembly [8], [9], [26], [27]. In contrast, Tpm1.8 is more abundant in lamellipodia where it promotes lamellipodial persistence and cell motility by inhibiting Arp2/3 activity in vivo [28].

In the past decade, a common approach to produce and characterize cytoplasmic Tpm isoforms has been to use constructs with a fused fluorescent protein. In this method, an Arg-Ser motif mimics the N-terminal acetylation required for Tpm function [8], [14], [29], [30], [31], [32]. However, given that Tpm interacts by their N- and C-termini, both pseudo-acetylation and fusion labelling disrupt normal cluster assembly [14]. Furthermore, in mammalian cells, the N-terminal methionine of short Tpm isoforms (here Tpm1.8 and Tpm3.1) is removed and the acetylation is carried by the second amino-acid [13].

Two methods have been proposed to bypass these difficulties but, to our knowledge, have never been combined. In order to produce a natural acetylation, we coexpress Tpm isoforms and the N-acetyltransferase NatA or NatB in BL21 *E. coli* [33]. For Tpm1.6 and Tpm2.1, the first methionine is directly acetylated by NatB. For Tpm1.8 and Tpm3.1, the first methionine is clipped by the bacterial methionine aminopeptidase MAP1, and the alanine in second position is acetylated by NatA. [13]. We verified by intact protein mass spectrometry that the purified Tpm1.6 and Tpm3.1 were fully modified (Fig. S1A).

For fluorescence labelling, we engineered cysteine mutants of each Tpm isoform [7]. All WT cysteines were replaced by lysines, and a single cysteine was added, on the third amino acid of exon 3, corresponding to positions 47 or 83 for short or long Tpm isoforms, respectively. This position was chosen as it does not interfere with Tpm dimerization and Tpm-actin interaction. Cysteine mutant Tpm were labelled by incubating with Alexa Fluor Maleimide (Method).

We first characterised the assembly of Tpm clusters onto actin filaments, one isoform at a time. To do so, we polymerized single actin filaments from surface-anchored spectrin-actin seeds, inside a microfluidic chamber [34]. Since Tpm bind weakly to fluorescently labelled actin, we assembled filaments with a long unlabelled stretch, flanked by two labelled actin segments (Fig. 1B, C) [32]. Since all studied isoforms are localised to non-sarcomeric structures in cells [6], we used β-actin, co-expressed in yeast *Komagataella pastoris* (formerly *Pichia pastoris*) with enzymes Naa80 and SETD2, to ensure actin N-terminal acetylation and His73 methylation, respectively [35]. Given that the second labelled actin segment was polymerized for at least 5 min, the unlabelled β-actin segment was in a fully (more than 99%) ADP-state [36]. We then exposed these filaments from time t=0 onwards to a constant concentration of fluorescently-labelled Tpm. Finally, we manually analysed on kymographs the nucleation and the elongation rate of Tpm clusters (Fig. 1E).

As previously described, we observed Tpm clusters to nucleate uniformly along the filament length (Fig S1B), and to elongate towards the two filament ends, for all isoforms (Fig. 1E). However, different isoforms appeared to bind with different dynamics: Tpm1.6 and Tpm1.8 tended to nucleate sparsely but to elongate rapidly, while Tpm2.1 and Tpm3.1 had the opposite behaviour and required higher concentrations to cover filaments.

We then measured the elongation speed of Tpm1.6, Tpm1.8 and Tpm2.1 clusters depending on the Tpm concentration. The elongation speed increases linearly with the Tpm concentration, in agreement with a simple kinetic model of Tpm dimer binding and unbinding with rate constants k_on_ and k_off_, respectively (Table 1).

**Table 1:**
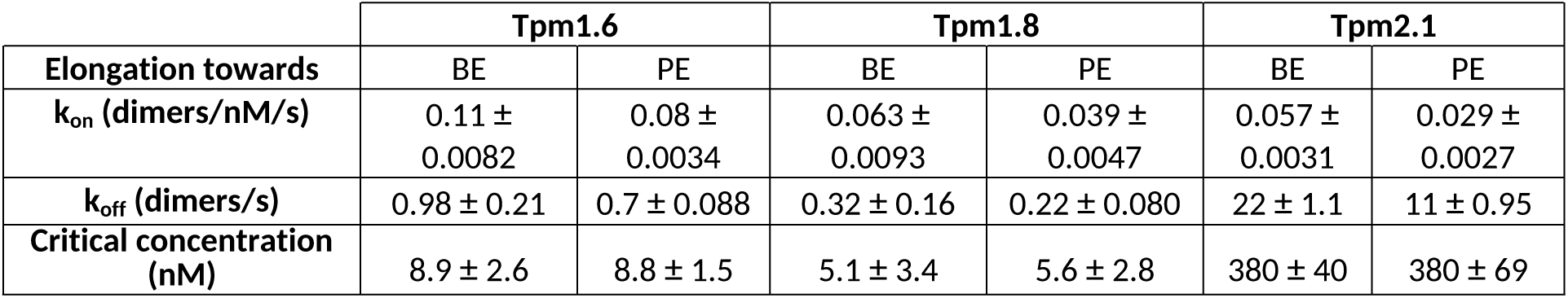
Kinetics of Tpm isoform cluster assembly and disassembly. Rate constants were obtained by fitting elongation and disassembly rates (Fig. 1F). Best fit values and standard deviations.

Consistent with thermodynamics, the critical concentration at the two Tpm cluster ends (towards the filament BE and PE) are equal (Fig. 1F, Table 1). Interestingly, we observed that, for the three isoforms, Tpm clusters are more dynamic (higher k_on_ and k_off_) at the cluster ends towards the BE of the filament. In order to exclude an effect of the flow, we performed control experiments with filaments anchored by their barbed end with Capping Protein (CP, Fig. S1C) and confirmed the biased elongation towards the barbed end. This observation is consistent with previous observations with fusion-tagged proteins [12] and seems to be conserved among all Tpm isoforms. Unfortunately, we could not perform a similar analysis with Tpm3.1 which offered a very limited range of concentrations at which we could measure cluster elongation because of the fast nucleation.

We then compared the relative contribution of nucleation vs elongation in the assembly of Tpm clusters. Because Tpm isoforms show vasty different binding dynamics, we first adjusted their concentrations such that they elongate at the same rate and then compared their nucleation rate (Methods, Fig. S1D). Consistently with previous preliminary observations, we found that Tpm1.6 was the slowest nucleator (relative to elongation), followed by Tpm1.8, Tpm2.1 and Tpm3.1 (Fig. 1G).

### 2. Tpm isoforms can coassemble to form mixed clusters

In a previous publication, Gateva et al. showed in vitro that Tpm isoforms from gene TPM1 cannot coassemble with isoforms from genes TPM2 or TPM3 [8]. This result was supported by the fact that Tpm1.6 and Tpm3.2 have different positions over actin filaments, showing a 5Å translation between the two isoforms, as revealed by cryoEM [37]. We wanted to revisit this observation taking advantage of our Tpm constructs and microfluidic setup which allows us to follow over time the coassembly or exclusion of Tpm clusters.

Using the same setup described above, we now exposed unlabelled β-actin segments from time t=0 to a solution containing two Tpm constructs labelled with Alexa Fluor 488 and Alexa Fluor 568 (Fig. 2A). Note that the two constructs could be of the same or different isoforms. Because the effective affinity highly varies between isoforms, we adjusted the Tpm concentrations such that: 1. we could observe single isolated clusters nucleating and assembling, and 2. the two isoforms appeared to have qualitatively the same fluorescence level over the filaments. For each actin filament, we then constructed a kymograph showing the assembly of Tpm clusters and measured the colocalisation coefficient “*r”* as the normalized product of the 488- and 568-nm fluorescence intensities (see Methods).

As positive controls, we measured the colocalisation coefficient for the same isoform labelled with two different colours (Fig. 2B). With the two controls, Tpm1.6 and Tpm3.1, we found a mean coefficient *r* between 0.6 and 0.8 (Fig. 2D). As a negative control, we assembled randomized kymographs whose colours were taken from different filaments, within the same experiment (Fig. 2B), and systematically found a mean colocalisation coefficient between 0.2 and 0.35 (Fig. 2D).

We then tested five different combinations of Tpm1.6, Tpm1.8, Tpm2.1 and Tpm3.1, and for all found a mean colocalisation coefficient above 0.5, while randomized kymographs had a mean coefficient below 0.4 (Fig. 2C-D), indicating that all isoforms can coassemble. Visually, we saw the fluorescence intensity varied spatially, with some pixels enriched in one isoform over the other. However, this aspect was equally present in control experiments (Tpm1.6/Tpm1.6 and Tpm3.1/Tpm3.1) suggesting that local variability could be attributed to stochastic incorporation of Alexa488- or Alexa568-labelled dimers.

Finally, we wondered whether Tpm coassembly might depend on the labelling strategy. However, we also observed sfGFP-AS-Tpm1.6 and mCherry-AS-Tpm3.2 to colocalise (Fig. S2B), in contradiction with previous observations [8]. In addition, we found our Tpm-Alexa constructs to coassemble in open chambers (Fig. S2A), in an experimental setup very similar to that of [8].

### 3. Tmod nucleates Tpm clusters at filament pointed end

In a cellular context, the nucleation and elongation of Tpm clusters are most likely regulated by other actin binding proteins. To our knowledge, all in vitro experiments have failed so far to identify proteins that would directly promote the assembly of Tpm clusters. Surprisingly, we saw that when actin filaments where anchored with spectrin-actin complexes, Tpm clusters would often appear from the anchor (Fig. S3A). We analysed the composition of our spectrin-actin complex by tandem mass spectrometry and identified tropomyosins and tropomodulin-1 (Tmod1), in agreement with a previous publication [22] (Supplementary data). We thus wondered whether tropomodulins could play the role of Tpm cluster nucleators. We compared two of the four human Tmod isoforms, Tmod1 and Tmod3, which are both widely expressed in different cell types.

To test whether Tmod can nucleate Tpm clusters at actin filament PE, we first performed experiments in open chambers (Fig. 3B). Pre-polymerized actin filaments were incubated with different Tpm isoforms (Tpm1.6, Tpm1.8, Tpm2.1 or Tpm3.1), in the presence or absence of Tmod (Tmod1 or Tmod3). In order to quantify the recruitment of Tpm, we measured the distance *d* between the first Tpm cluster that nucleated and the closest end of the filament, of length *l*, for each filament (Methods). We then calculated, for each experiment, the cumulative distribution function (CDF) of *d/l* (Fig. 3C).

As a first example, we compare the localisation of Tpm1.6 clusters, with and without Tmod 1 and Tmod3. In the absence of Tmod, Tpm clusters appear randomly along the length of the filaments: the cumulative distribution of the localisation of Tpm clusters along filaments (CDF) increases linearly from *d/l=0* to 0.5 (Fig. 3C). In contrast, in the presence of Tmod1 or Tmod3, the CDF increases sharply between *d/l=0* and *0.1*, i.e. many Tpm1.6 clusters appear to nucleate near one of the two filament ends (Fig. 3C).

In order to better quantify the recruitment of Tpm at the filament end, we record the value of the CDF at *d/l=0.04*, corresponding to approximatively 2 pixels from the filament end. If the spatial distribution is homogeneous, this value should be around 0.08. Higher values indicate a biased recruitment at the filament end. In the example above, the fraction of Tpm clusters near the filament end is around 0.1 without Tmod, 0.65 with Tmod1 and 0.85 with Tmod3 (Fig. 3D).

We next wondered whether this recruitment was Tpm- or Tmod-isoform specific. We thus tested all combinations between Tpm1.6, Tpm1.8, Tpm2.1 and Tpm3.1, and Tmod1 and Tmod3 (Fig. 3D). We found that for all combinations of Tpm and Tmod isoforms, the fraction of Tpm at filament ends was higher than 0.4. Interestingly, the values with Tmod1 were systematically smaller (around 0.5) than those with Tmod3 (around 0.75). This difference might either come from a difference in the affinity of Tmod for the PE, or from the efficiency of Tmod isoforms to nucleate Tpm clusters.

Since Tmod are pointed end cappers, we expect the Tpm clusters to be nucleated at the same filament end. However, in the experiments above, we cannot distinguish the two ends. In order to confirm that Tpm clusters are nucleated at the filament pointed end, we performed similar experiments inside a microfluidic chamber with filaments anchored at their BE by Capping Protein (Methods), leaving the PE free for Tmod to bind (Fig. 3E, F). We then study the nucleation of Tpm clusters at the PE by quantifying the position of the first Tpm cluster for each filament, as in open chamber experiments.

In the absence of Tmod, Tpm clusters appear to nucleate uniformly along the filament length (linear CDF, Fig. S3C). In contrast, when also exposing to Tmod1 or Tmod3, Tpm clusters readily appeared at the free pointed end (Fig. 3G, S3C). We further quantified the fraction of Tpm clusters appearing at the pointed end. Note that for all movies, we systematically analysed a 5.9 µm-long segment (45 pixels), starting from the free end. We confirmed with Tpm1.6 and Tpm1.8 that Tmod nucleates Tpm clusters at the actin filament PE (Fig. 3D, diamond symbols). As a control, we performed the same experiment with filament anchored with spectrin-actin seed and saw no direct recruitment of Tpm at the free BE, as expected (Fig. S3C).

In microfluidics, based on fluorescence level, we also noticed that Tmod did not nucleate the two Tpm clusters on the two filament strands simultaneously, but instead, one at a time (Fig. 3G). This suggest that Tpm-binding domains of Tmod nucleate Tpm clusters independently, likely with different efficiencies. We cannot exclude the possibility that Tmod detaches from the PE. Unfortunately, we tried to repeat the experiment with fluorescently labelled Tmod but were not able to observe individual fluorescent Tmod bound to actin, either because of poor labelling, challenges in imaging single molecules or rapid unbinding from actin PE.

### 4. Lmod nucleates Tpm clusters along the length of actin filaments

We thus tested whether Lmod, which are closely related to Tmod, could nucleate Tpm clusters. As a model, we used Lmod2 which is expressed primarily in cardiac muscle cells. We first exposed CP-anchored actin filaments to a solution containing Tpm1.6, with or without Lmod2, inside a microfluidic chamber. The addition of Lmod2 strikingly increased the nucleation of Tpm clusters, all along the filament (Fig. 4B). To quantify this effect, we measured the density of Tpm clusters over time and found a ∼5 fold increase in the nucleation rate of Tpm1.6 by Lmod2 (Fig. 4D-E).

Based on fluorescence intensity, each Lmod2 appears to nucleate a single Tpm cluster (Fig. 4B-C), consistent with the fact that Lmod only contains one Tpm binding site. In addition, Tpm clusters appeared to elongate towards the anchored BE only, with the PE side of the clusters remaining static over time. To confirm this directionality, we repeated the experiment with actin filaments polymerized from spectrin-actin seeds. Again, Lmod-nucleated Tpm clusters only elongated towards the BE (Fig. 4C). Lmod2 most likely remains bound to F-actin and acts as a Tpm cluster capper.

We then wondered whether Lmod2 would show specificity towards some Tpm isoforms and repeated the experiments with Tpm1.8, Tpm2.1 and Tpm3.1. Lmod2 showed similar effects with a higher nucleation rate for all Tpm isoforms, with no obvious preference for a Tpm isoform (Fig. 4E and S4).

## Discussion

Understanding the dynamics and regulation of Tpm cluster assembly over actin filaments has been a challenge for the field due to two primary obstacles: purifying functional Tpm and visualizing their binding. In this study, we propose a novel fluorescent constructs with comparable affinity to recombinant Tpms expressed in human Expi293F cells [13]. It will be important for future studies to examine other PTMs that can be modifying Tpm, such as phosphorylations and acetylations [13] which likely affect their regulation [38], [39].

A standard method to assess Tpm affinity involves cosedimentation assays with F-actin at various Tpm concentrations. By fitting the density of pelleted Tpm with a Hill function, one obtains the apparent dissociation constant (K_app_) and the Hill coefficient, which reflects cooperativity. However, since these measurements are conducted in bulk and at steady state, it is impossible to predict the dynamics and relative contributions of Tpm cluster nucleation versus elongation. Our method offers a novel perspective by examining individual Tpm cluster formation, allowing for better insight into differences in Tpm isoform binding dynamics. We discovered that Tpm1.6 and Tpm1.8 nucleate relatively few clusters that elongate rapidly, while Tpm2.1 and Tpm3.1 exhibit a strong nucleation rate relative to their elongation. These distinctions might lead to preferences for different actin filament populations in cells, depending on factors such as filament length or lifetime.

We also demonstrated that the four Tpm isoforms coassemble to form mixed clusters, in apparent contradiction with a previous report [8]. Despite our efforts in open chambers and with GFP/mCherry-fusion Tpm constructs, we never observed clear segregation between two Tpm isoforms. Nevertheless, it is still possible that under certain experimental conditions (such as protein concentrations and isoforms, additional actin binding proteins) Tpm isoforms could segregate into different clusters. In addition, we cannot rule out the possibility that Tpm isoforms form microdomains or islands within clusters, below the optical resolution, which could be enriched in one isoform versus another.

Finally, we found that Tmod and Lmod nucleate Tpm clusters at the pointed end and along the length of actin filaments, respectively. These findings raise a number of questions on the molecular mechanisms leading to Tpm cluster nucleation, and underscore the need for further experiments. Specifically, it remains unclear whether Tpm and Tmod/Lmod form a complex in solution or if Tmod/Lmod binds first to F-actin, recruiting Tpm which then stabilizes Tmod/Lmod. Additionally, understanding how Lmod binds along F-actin is crucial. We recently reported, in cryoEM, to only observe Lmod at filament pointed end, despite the presence of Tpm1.1 [24]. These structures reveal that Lmod’s Leucine Rich Region (LRR) modifies the conformation of the last actin subunits at the filament PE but appears to be unable to bind along F-actin [24]. Nevertheless, the fact that Lmod-nucleated Tpm clusters are permanently capped at their PE side indicate that Lmod remains strongly bound, probably to both the filament side and Tpm. Additional interaction with Tpm and with F-actin through Lmod’s WH2 and proline rich domains likely stabilise Lmod along the filament length.

To our knowledge, Tmod and Lmod are the first proteins reported to nucleate Tpm clusters. Formins have long been considered potential candidates for this role [16], and formin DAAM1, in U2OS cells, has recently been shown to assemble actin networks specifically enriched in Tpm3.2 [40]. However, despite our efforts and those of other labs [15], [17], we have not been able to detect direct recruitment of Tpm by formins mDia1 or DAAM1. One strong limitation of these in vitro reconstitution experiments is that the formin constructs used comprise only the FHL1-FH2-DAD domains, lacking the N-terminal regulatory domains that might interact with Tpm. It is also plausible that formins mediate the recruitment of Tpm indirectly, possibly with Tmod/Lmod acting as intermediates. Other formin partners might also be involved, particularly in yeast, which lack Tmod/Lmod. Furthermore, Tpm assembly might be indirectly regulated by competition with other actin-binding proteins, in the presence of different actin architectures [15]. Future in vitro reconstitution experiments will be essential to elucidate the full regulatory mechanism of Tpm isoform assembly by actin regulators.

## Acknowledgments

H.W. was supported by the Agence Nationale de la Recherche ANR-21-CE11-0006–01. A.J. Was supported by the Impulscience program from the Bettencourt-Schueller Foundation (Grant #1235). C.B. was supported from the investment program “France 2030” as part of the IdEx program (ANR-18-IDEX-0001) implemented by Université Paris Cité, under which the inIdEx project Formula is conducted. R.D. was supported by National Institutes of Health Grant R01 AR087343. MS analysis at ProteoSeine facility were financially supported by Région Ile de France (SESAME 2021 EX061040), CNRS and Paris Cité University.

## Data availability

Summary of the tandem mass spectrometry analysis of spectrin-actin complexes: only proteins with a total area larger than 10^5^ and area of unique peptides larger than 10^3^ are listed.

## Methods

### Protein production and purification

Rabbit skeletal muscle actin (UniProt P68135) was purified from acetonic powder, following the protocol from [41] as described in [42]. Rabbit actin was fluorescently labelled on accessible lysines with sulfo-NHS-Alexa568, following [42]. Recombinant human β-actin (UniProt P60709) was produced in Komagataella pastoris (formerly Pichia pastoris) following the protocol by Hatano et al. [35]. Note that β-actin was coexpressed with NAA80 and SETD3 to ensure proper N-terminal acetylation and His73 methylation, respectively.

Spectrin-actin seeds from human red blood cells were purified as described in [42]. Recombinant human profilin1 (UniProt P07737) and chicken Capping Proteir CapZ α1-β1 with His and SNAP tags on the β1 subunit (Uniprot: P13127 and P14315) were expressed and purified as described in [42]. Recombinant human tropomodulin1 and leiomodin2 were purified as described in [43]. Recombinant human sfGFP-AS-Tpm1.6 and mCherry-AS-Tpm3.2 were purified as described in [32].

Mouse Tropomodulin3 (Uniprot Q9JHJ0) was expressed in BL21 cells and induced at 37°C for 2h. Cells were lysed by sonication and GST-tagged Tmod3 was captured over GST-beads (1h, 4°C) and eluted with 20 mM Tris-HCl pH 8.0, 150 mM NaCl and 2.5 mM CaCl2. The tag was cleaved by adding thrombin (1 unit per 100 µg of protein, 30 min, 30°C). The reaction was stopped by adding 2 mM PMSF. The protein was dialysed overnight in 20 mM Tris-HCl pH 8.0, 150 mM NaCl, 1 mM DTT and further purified over a gel filtration column HiLoad 16/600 Superdex 200. Protein concentration was determined with a BCA assay, flash frozen in liquid nitrogen and stored at −70°C.

In order to label Tpm isoforms with maleimide-Alexa488 or maleimide-Alexa568, we mutated all cysteines to lysines, and changed one alanine to cysteine such that labelling does not disrupt Tpm-Tpm, Tpm-actin and Tpm-Tmod/Lmod interactions. The mutants were as follows: Tpm1.6 C193L A83C, Tpm1.8 C157L A47C, Tpm2.1 A83C, Tpm3.1 C154L C170L C226L A47C. Note that because Tpm1.8 and Tpm3.1 are shorter isoforms that lack the second exon, amino acid A47 in Tpm1.8 and Tpm3.1 correspond in sequence alignment to A83 in Tpm1.6 and Tpm2.1. Tpm were coexpressed in BL21 cells with NatA (Tpm1.8 and Tpm3.1) or NatB (Tpm1.6 and Tpm2.1) to ensure N-terminal acetylation and induced overnight at 16°C [10]. We confirmed by intact protein mass spectrometry that virtually all Tpm monomers were acetylated (Fig. S1A). E. coli were lysed by sonication in lysis buffer (20 mM Sodium Phosphate pH 7.5, 500 mM NaCl, 5 mM MgCl2, 1 mM DTT, protease inhibitor and DNAse I) and were heated to 90°C for 10 min. Debris were removed by centrifugation (100 000 g, 45 min). We then performed a pH drop (bringing the pH to 4.5 with HCl) and further purified Tpm on a CHT Type I column, eluted with a linear gradient of Sodium Phosphate from 10 to 200 mM, along with 1 M NaCl and 0.5 mM DTT. Protein concentration was determined with a BCA assay. For labelling, Tpm were first dialysed overnight in 20 mM TrisHCl (pH = 7.0), 100 mM KCl, 5 mM MgCl2. The next day, maleimide-Alexa488 or maleimide-Alexa568 was added with a 2x excess and left for 2h at room temperature on a wheel. The reaction was stopped by adding 0.5 mM DTT and protein were finally dialysed overnight in 20 mM TrisHCl (pH 7.0), 100 mM KCl, 5 mM MgCl2, 0.5 mM DTT.

All experiments were performed in F-buffer: 10 mM Tris-HCl pH 7.4, 100 mM KCl, 1 mM MgCl2, 0.2 mM EGTA, 0.2 mM ATP, 10 mM DTT, 2 mM DABCO.

### Intact protein mass spectrometry

Tpm samples (500 ng) were injected onto a custom made BioResolve RP (450Å, 2.7 μm, 0.3 mm x 150 mm) Polyphenyl Column (Waters, Milford, MA, USA) using an Acquity M-Class chromatographic system from Waters. Mobile phases A and B were H2O and H2O/ACN 20/80, respectively, acidified with 0.1% (v/v) DFA. Samples were eluted for 70 min using the percentage of acetonitrile described in the table below. The eluent was sprayed using the conventional ESI ion source of a Waters Cyclic IMS SELECT SERIES mass spectrometer operating in the positive ion mode. The quadrupole was operated in RF-only mode, the TOF was operated in V-mode with a resolution specification of 60,000 FWHM at a mass range of m/z 50-4000. Data were deconvoluted using the Intact Mass v5.3.44-gbc4a1db565 software provided by Protein Metrics (Fig. S1A).

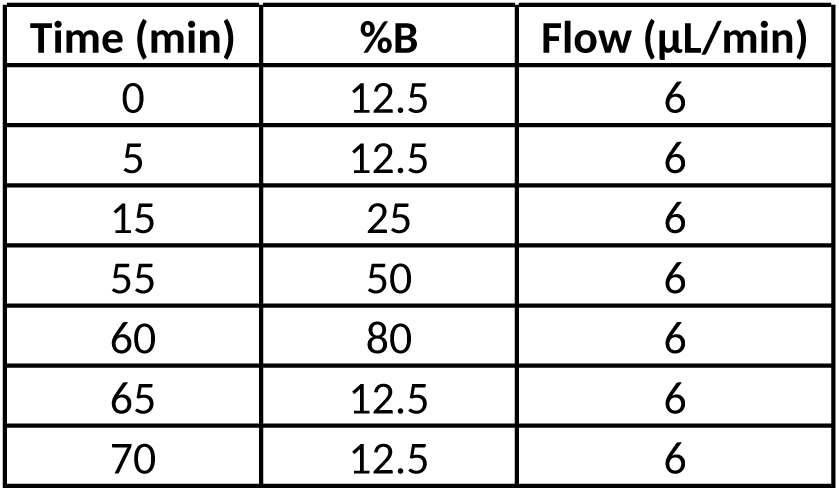

### Microfluidic chamber and single filament reconstitution

Single actin filaments were reconstituted inside a microfluidic chamber as in [32]. The microfluidic chambers were made of Poly Dimethyl Siloxane (PDMS, Sylgard). The channels were cross-shaped with 3 inlets and 1 outlet, 23 μm high and 800 μm wide. The PDMS chambers are mounted on standard glass coverslips (VWR, #1.5). To do so, glass coverslips are first cleaned with 2% glass cleaning solution (Hellmanex™ III), 2M KOH, dH2O and ethanol, under 30 min sonication for each step. When starting an experiment, a microfluidic chamber was prepared by exposing a PDMS chamber and a cleaned glass coverslip to air plasma for 30 s and then binding the two together [34].

Microfluidic chambers were connected to solutions containing proteins and injected by applying pressure in the gas phase using the software Oxygen (Fluigent). Full description of the microfluidic device and typical usage can be found in [34]. Briefly, inlets were filled with solutions containing, for example, buffer only, G- or F-actin, Tpm isoforms, supplemented with Tmod or Lmod. The field of view was sequentially exposed to these solution, changing from one solution to another in less than a second. While only the applied pressure was imposed, flow meters allowed us to ensure the correct flowing of solutions. Note that in these micro chamber, solutions do not mix.

### Pointed-end anchoring with Spectrin-actin seeds

For the experiments detailed in sections 1 and 2, 20 µL of a solution of 11.7 pM spectrin-actin seeds was injected in the chamber with a pipet for seeds to adhere non-specifically to the glass surface. After 1-2 min of incubation, the chamber was passivated overnight by injecting 20 µL of 0.1 mg/mL PLL-PEG (SUSOS) in 10 mM HEPES pH 7.4 with a pipet. The next day, the chamber was rinsed by injecting 400 µL of F-buffer with a pipet and then connected to the microfluidic system (Fluigent MCFS and Flow Units).

Actin filaments were then polymerized by exposing alternatingly to labelled and unlabelled G-actin solutions in order to obtain filaments containing two short fluorescent segments that reveal the two ends, and a long central unlabelled segment to which Tpm bind. First, short labelled actin segments were polymerized by flowing in 0.5 μM 10% Alexa Fluor 568–labeled α-skeletal G-actin for 5 min. Then, long unlabelled actin segments were polymerized by flowing in a solution with 1.2 μM cytoplasmic β G-actin and 1.3 µM profilin-1 (to avoid actin spontaneous nucleation) for 15 min. The solution of 0.5 μM 10% Alexa Fluor 568–labeled α-skeletal G-actin was flowed again for 5 min.

From time *t=0* onwards, the actin filaments were exposed to a solution containing tropomyosin to assemble Tpm clusters.

### Barbed end anchoring with Capping Proteins

Biotin-SNAP-CapZ was first prepared by incubating SNAPtag-CapZ with SNAP-biotin (equimolar concentration), supplemented with 1 mM DTT, for ∼1h at RT. Actin filaments were prepolymerized at 4 µM by mixing unlabelled β-actin and Atto642-labelled α-skeletal-actin (final labelling 5%, about 10% α-skeletal-actin).

The microfluidic chamber was passivated by injecting 20 µL PLL-PEG 1% biotin (SUSOS) in 10 mM HEPES, with a pipet. The solution was left >1h at RT. The chamber was then connected to the microfluidic controller (Fluigent) and thoroughly rinced with 500 µL F-buffer. The chamber was then functionnalized by exposing the surface to 400 nM biotin-SNAP-CapZ (5 min). Actin filaments (diluted down to 0.5 µM) were injected at a slow flow (<500 nL/min), captured at their barbed end by CapZ. Finally, filaments where exposed from time *t=0* onwards to Tpm with or without Tmod/Lmod.

### Data acquisition

Observations were made using a Nikon Ti2 inverted microscope equipped with a 100× oil-immersion objective and a Kinetix 22 camera. The temperature of the microfluidics chamber was maintained around 25°C using a collar objective heater (Okolab). We used an azimuthal total internal reflection fluorescence (TIRF) illumination setup (iLAS2, Gataca Systems), with 488-, 561-, and 642-nm tunable lasers (maximum power 100 mW each). The manipulation and adjustment of the TIRF setup were carried out using MicroManager software (version 2.0.1).

The Tpm clusters nucleation and elongation were imaged with an acquisition rate between 1 frame every 500 ms to 1 frame every 3 s depending on the nucleation/elongation speed of the Tpm clusters. The acquisition was started a few seconds before exposing filaments to Tpm (*t=0* as defined above). Nucleation and elongation rates were later measured as described below.

### Nucleation and elongation rates analysis

Image analysis was performed manually on Fiji. First, we excluded filaments that either did not contain the two labelled segments clearly visible, or were sticking to the surface. We then constructed kymographs of all other filaments.

To quantify the nucleation rate, the frame and position of nucleation for every Tpm cluster was manually recorded from these kymographs. Elongation rates were measured for each cluster by manually tracing the slope on the kymographs (Fig. 1E).

Data were pooled, fitted and plotted in Python (Jupyter).

### Coassembly analysis

As previously, we only analysed filaments with clearly visible, non-sticking ends. Then, for each filament, we constructed a two-colour kymograph corresponding to the two Tpm constructs. The kymographs only started when filaments were exposed to Tpm and ended when clusters started to overlap (typically around 5 clusters per filament).

For each kymograph and colour, we subtracted the background, estimated as the modal fluorescence value of the image, so that every pixel with no Tpm bound has an intensity close to 0 (negative values were set to 0), yielding the net fluorescence intensities *I_488_* and *I_568_*. We then computed for each filament the correlation coefficient as 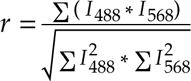 where the sums are taken over all pixels of the kymograph.

As the background has been subtracted, if the two Tpm construct never coassemble, i.e. *I_488_* or *I_568_ = 0* on all pixels, then *r = 0* On the other hand, if the two Tpm constructs perfectly colocalise, i.e. *I_488_* = *I_568_*, then *r = 1*.

The same procedure was applied for the negative controls (“randomized” kymographs). However, we assemble kymographs whose colours were taken from two different filaments: the 488 channel from filament *n* and 568 channel from filament *n+1*.

Colocalisation coefficients were plotted using the web app SuperPlotsOfData [44].

### Localisation of Tpm nucleation by Tmod

In order to determine whether Tmod could specifically nucleate Tpm cluster at the filament pointed end, we first adjusted the concentration of the Tpm such that few clusters would appear over a couple of minutes.

### Open chambers

Chambers were assembled with two ethanol-cleaned and dried coverslips (VWR, #1.5), and separated by thin strips of double-sided tape (Fig. 3B), typically making 4 channels (about 5 µL each). Surface were passivated with 2.5% BSA in F-buffer (5 min at RT). 100 nM preassembled actin filaments (β-actin labelled with 5% Atto642-α-skeletal-actin) were mixed with Tpm, with or without 50 nM Tmod, in F-buffer supplemented with 0.25% methylcellulose, and rapidly injected into the chamber (<1 min before the start of the acquisition).

Filaments and Tpm were imaged, typically one frame every second in order to visualise the appearance of the first Tpm clusters. To analyse the localisation of Tpm nucleation, we selected in Fiji for filaments with a standard length (typically between 2.5 and 10 µm, average ∼7 µm) for which the nucleation of the first Tpm cluster was clearly visible. We then manually measured the filament length, *l*, and the distance between the center of the Tpm cluster and the closest filament end, *d* (as a consequence *d*≤*l/2*). Again, only the first Tpm nucleation event was noted for each filament. For each movie we then computed the cummulative distribution function of *d/l* between 0 and 0.5. For a given relative distance *d/l\**, the function corresponds to the fraction of clusters with *d/l≤d/l\**. If the distribution of nucleation event is homogeneously distributed along the filament length, the cummulative distribution function is then linear. If all nucleation event occur at either filament end (*d=0*), then the function sharply increases at small values.

We then defined the fraction of Tpm clusters near the filament end as the value of the cummulative distribution function at d/l = 0.04. For a perfectly uniform distribution, the fraction should be 8%.

### Microfluidic chambers

Single filaments were reconstituted as described above, either attached with spectrin-actin seeds (free barbed end) or CapZ (free pointed end), always with β-actin supplemented with Atto642-α-skeletal-actin (5% final labelling fraction) for visualisation. Filaments were then exposed to 20 nM Tpm1.6 or Tpm1.8 and imaged.

Movies were analysed as in open chambers, with the exception that we only analysed *l=45* pixel-long (5.9 µM) segments and that *d* was always measured from the free end. As a consequence *0≤d≤l*. Filaments shorter than 45 pixels were excluded from the analysis.

## Supplementary figures

**Figure S1:**
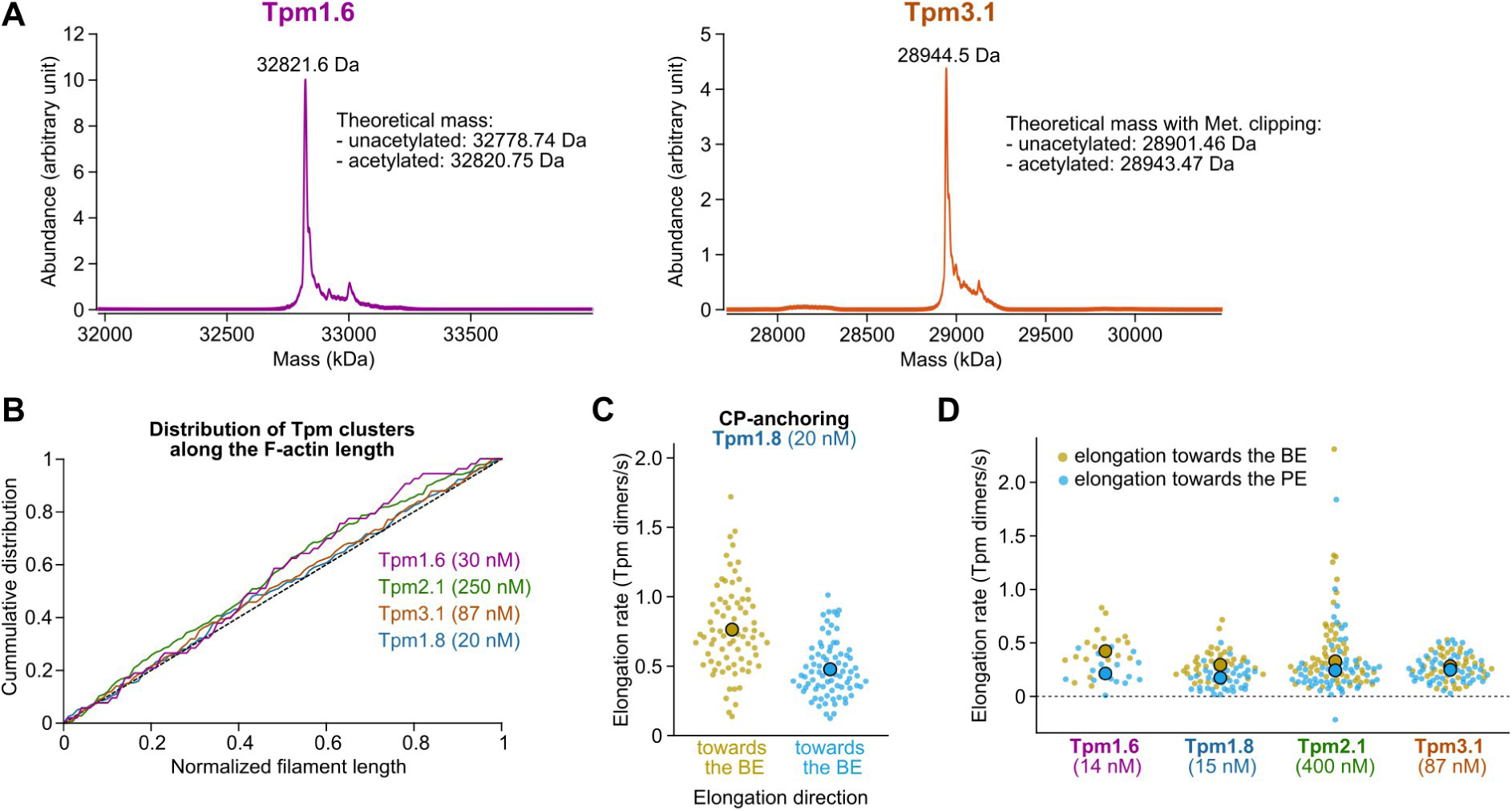
Characterisation of Tpm isoform and their binding to actin filaments. **A.** Mass spectrometry analysis of intact Tpm1.6 and Tpm3.1 confirm their N-terminal acetylation. The unacetylated form is not detectable. **B.** Homogeneous distribution of Tpm cluster along the actin filament length. The length is normalized from one end to the other of the unlabelled actin segment. The linear dashed line is shown as a reference for a purely uniform distribution. **C.** Biased elongation is also observed when filaments are anchored by their BE with CP-anchoring. **D.** Tpm isoform concentrations were adjusted such that their elongation was nearly equal, in order to compare their relative nucleation (Fig. 1G). C-D: small dots corresponds to single cluster measurements, large dots: averages over the experiment.

**Figure S2:**
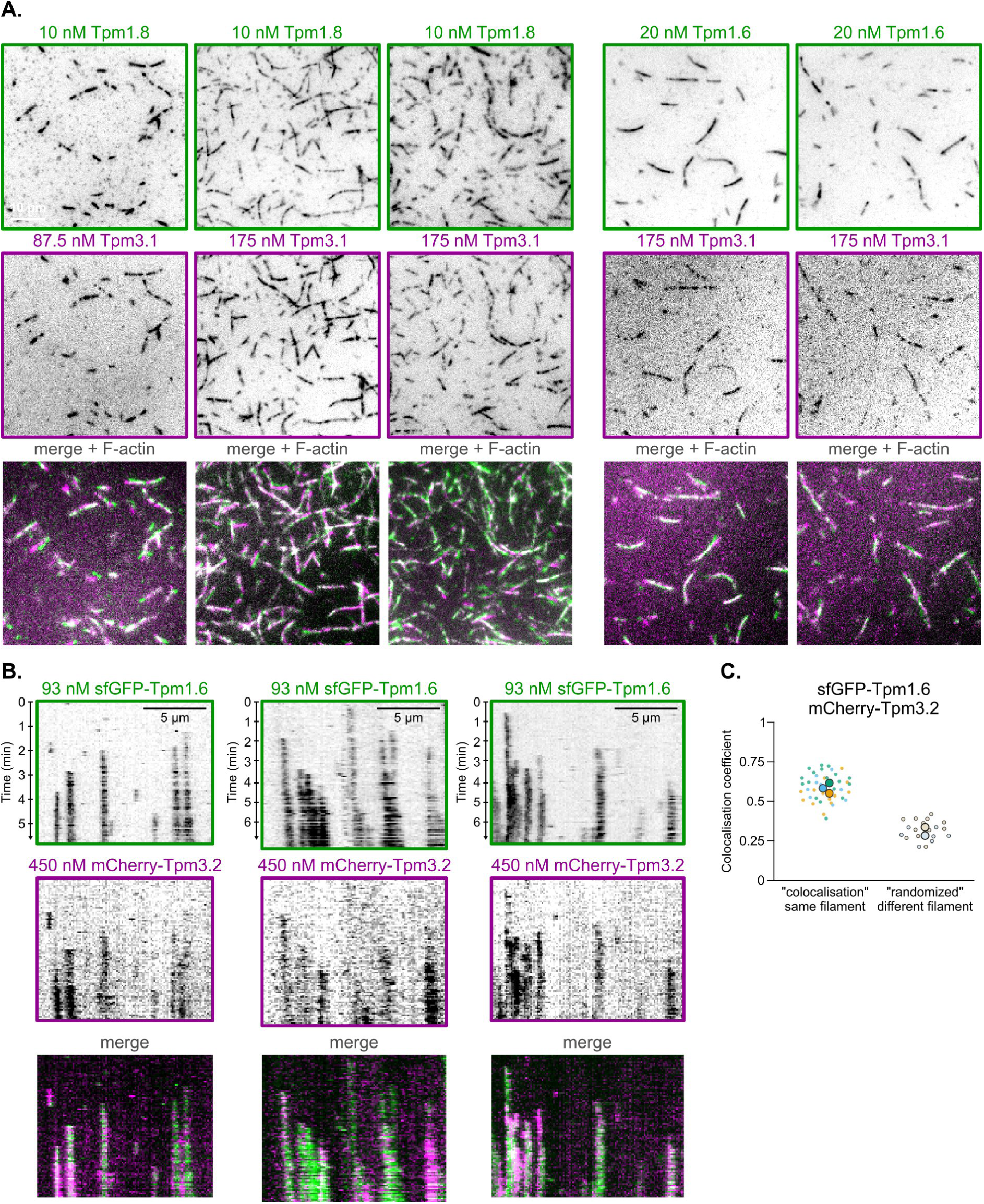
Tpm isoform co-assembly controls. **A.** Tpm1.6 and Tpm1.8 coassemble with Tpm3.1 in open chambers. Pre-assembled actin filaments were incubated with Tpm in F-buffer supplemented with 0.25% methylcellulose and rapidly injected between two glass coverslips separated with double-sided tape. Note that, since filaments are not attached to the surface, they diffuse such that the two colours might not appear in perfect register. **B.** sfGFP-Tpm1.6 and mCherry-Tpm3.2 coassemble in microfluidics. Typical kymograph showing the binding of the two Tpm isoforms (same constructs as in [8]) to unlabelled actin filaments. **C.** Colocalisation coefficient of sfGFP-Tpm1.6 with mCherry-Tpm3.2. Small dots corresponds to single kymographs, large ones to averages over single experiments.

**Figure S3:**
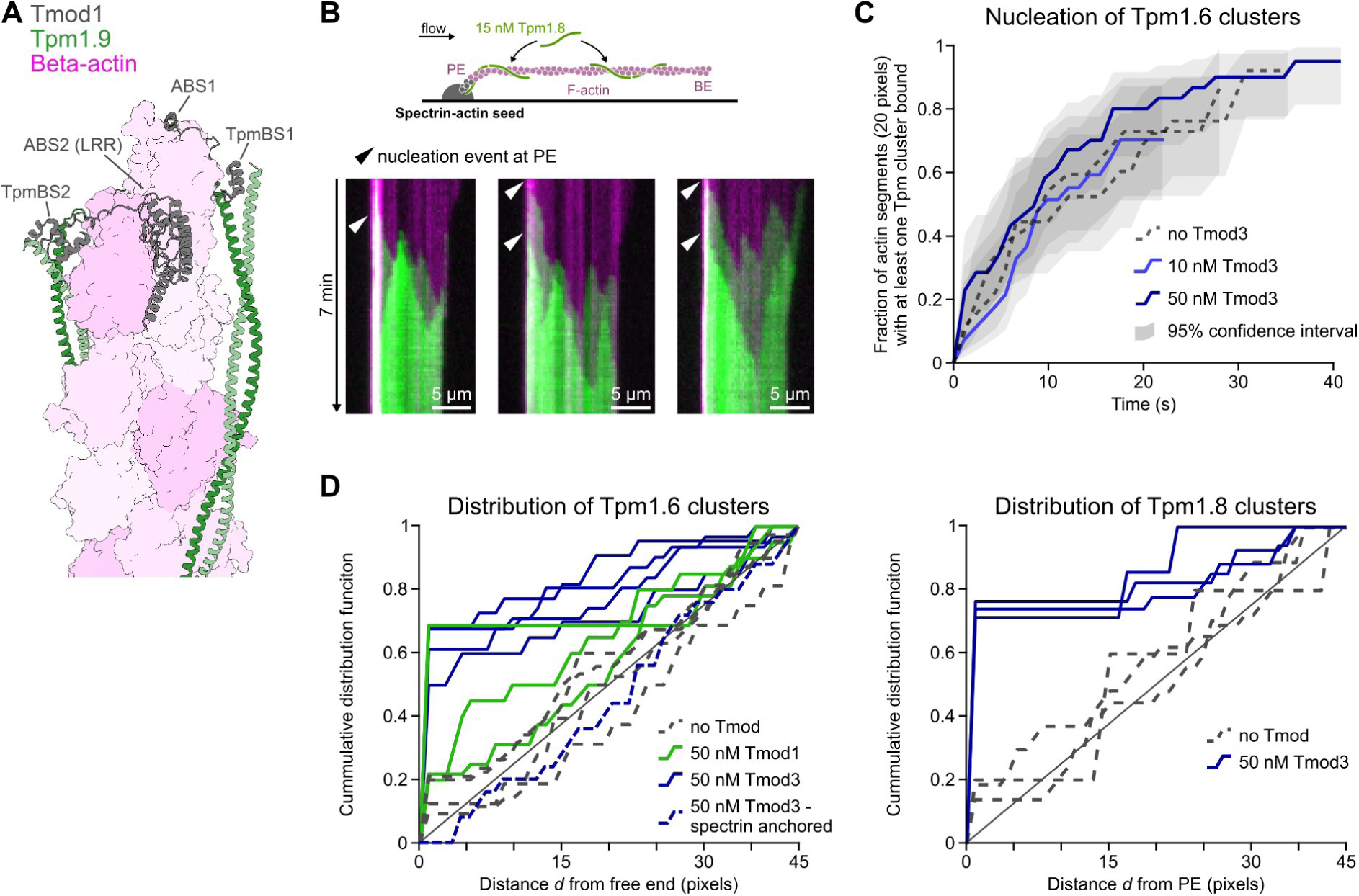
Spectrin-actin seeds and Tmod nucleate Tpm clusters at filament PE only. **A.** Structure of a pointed end (pink) decorated by a Tmod1 (dark grey) and two Tpm1.9 dimers (green). Adapted from [45], PDB IAH. **B.** Tpm1.8 clusters nucleate near the surface-anchored spectrin actin seed. Top: sketch of the experiment. Bottom: typical kymograph showing nucleation events near the seed (white arrow head). **C.** Quantification of Tpm1.6 nucleation along filament length, with or without Tmod3. Filaments were divided in 20 pixel-long segment and, for each segment, we noted the frame at which the first Tpm cluster appeared. Plotted are the fraction of cluster with at least one Tpm cluster i.e. (1 – survival function). Curves and confidence intervals were calculated with the Kaplan Meier method. N = 10-15 filaments, 39-55 clusters. Related to Fig. 3E-G. **D.** Distribution of Tpm cluster nucleation in microfluidics. Only the first nucleation was analysed, along a 45 pixel-long segment starting at the free end. All experiments were performed with Capping Protein-anchored filaments (free PE), except for the dashed blue line (left, “spectrin-anchored”, free BE). Note that for the later experiments, the spectrin seeds were more than 45 pixels away from the free end and, thus, nucleation event at the seed were not taken into account. N = 16-43 filaments (Tpm1.6) and 5-29 (Tpm1.8).

**Figure S4:**
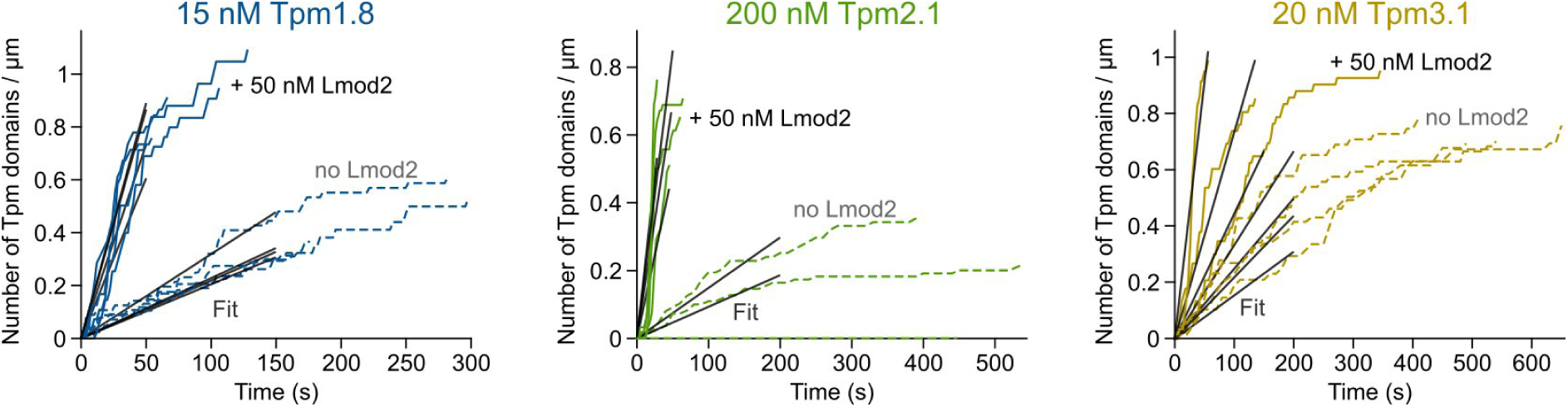
Nucleation of Tpm clusters by Lmod2. Quantification of the density of Tpm1.8, Tpm2.1 and Tpm3.1 clusters over time, with (solid lines) or without Lmod2 (dashed lines). Each line corresponds to a single experiment. Tpm1.8: N = 4-10 filaments, 18-42 clusters. Tpm2.1: N = 4-10 filaments, 24-43 clusters (and one movie without any cluster nucleating). Tpm3.1: N = 5-8 filaments, 36-56 clusters. Linear fits (grey lines) were used to calculate the nucleation rate (Fig. 4E).

